# Configural and featural processing, and their role in the face network

**DOI:** 10.64898/2026.09.04.749546

**Authors:** Yuxuan Zeng, Ren Hentz, David Osher

## Abstract

Object recognition can rely on the detection of individual features, as well as the spatial configuration of features, which is especially prominent in the identification of faces. We previously showed that canonical ‘face effects’ (e.g., inversion, part-whole) can be elicited by non-face stimuli that are identified strictly by configuration, but not in stimuli that are identified by features. Here we asked 1) whether there exist distinct neural bases that support domain-general configural vs. featural processing; and 2) how domain-specific face-areas may respond and interact with these processors. We trained adults to recognize these configural and featural stimuli, and then used fMRI to observe neural responses associated with each type of processing. Configural stimuli drove responses in superior parietal, superior frontal, and lateral occipitotemporal cortices, while featural stimuli primarily elicited responses in the ventral fusiform gyrus. Next, we identified face-areas with a separate fMRI localizer; the Occipital Face Area (previously linked to the identification of face parts) was selectively activated during recognition of featural stimuli, while the Fusiform Face Area was ambivalent, preferring neither configural nor featural stimuli. The posterior Superior Temporal Sulcus face-area, typically involved in the moveable aspects of faces (e.g. facial expressions), selectively responded to configural stimuli over featural stimuli. Finally, we observed that the functional connectivity of these face-areas to configural or featural regions mirrored this functional pattern. These domain-general processors and their differential connectivity with the face network may offer a mechanism by which face-areas use this information to accomplish their distinct roles in face perception.

**Significance Statement:** Object recognition can rely on the detection of individual features of a stimulus, as well as the spatial configuration of features, which is especially prominent in the identification of faces. However, holistic effects can also be observed for non-face abstract stimuli that have been trained to be recognized by only their configural or featural properties. Here, we characterize the neural systems involved in this domain-general configural vs. featural processing of abstract stimuli and how they interact with different parts of the face network, as a means to better understand the mechanisms of face perception.

## Introduction

To recognize an object, the visual system can rely on identification of individual features as well as the spatial configuration of features. Configural processing lies at the heart of face perception (Maurer et al., 2002; Tanaka & Farah, 1993; Young et al., 1987), and has long been treated as special to faces, or at most to categories of visual expertise (Farah et al., 1998; Gauthier & Tarr, 1997; McKone et al., 2007). However, it is also possible that configural processing is not a specific property for faces, but is instead recruited whenever recognition must draw on spatial relations. We recently found direct support for this idea. Abstract shapes that resemble no familiar object, categorizable only from the spatial relations among their parts, elicited the same hallmarks of holistic processing that mark face perception, such as the inversion effect and the parts effect (Zeng et al., 2026). These findings suggest that extraction of visual features and their spatial configuration can be domain-general rather than specific to face processing. Here we ask whether these domain-general processes are supported by distinct neural systems, and how they may facilitate face perception by interacting with the domain-specific face network.

Ventral occipitotemporal cortex is organized from perceptual to categorical representations along its posterior to anterior axis (DiCarlo et al., 2012; Grill-Spector & Weiner, 2014). Featural information is available before a stimulus is categorized, so it should be read out before the categorical end of that axis. Configural information instead requires relating parts in space, and such relations are represented at several stages of the hierarchy rather than at one place on it, leaving open where a configural preference should fall. For faces, the two types of processing have separate neural bases (Zachariou et al., 2017), but whether that holds once category-specific information is removed (as in the case of abstract, novel stimuli) is unknown.

A second question concerns the network that reads this information out. Face recognition is supported by a set of regions including the fusiform face area (FFA), occipital face area (OFA), and posterior superior temporal sulcus (pSTS); these are among the most well-characterized categorical regions in visual cortex (Haxby et al., 2000; Kanwisher et al., 1997; Puce et al., 1998). Yet it still remains unknown how each region contributes to face recognition. Evidence from lesion, adaptation, and holistic-processing studies suggest that the FFA is critical for face perception (Barton et al., 2002; Schiltz & Rossion, 2006; Yovel & Kanwisher, 2005; Zhang et al., 2012) while the pSTS supports recognition of facial expressions or dynamic aspects of faces (Allison et al., 2000; Puce et al., 1998, 2003); both FFA and pSTS may thus favor configural processing, while the OFA, placed early in the hierarchy as a processor of local parts, may be more involved in the featural processing of faces (Arcurio et al., 2012; F. Nichols, 2010; Pitcher et al., 2007). Yet each region has also been reported with the opposite bias (Bona et al., 2016; Flack et al., 2015; Golarai et al., 2015; Harris & Aguirre, 2008; Kho et al., 2023; Liu et al., 2010; Renzi et al., 2015; Rhodes et al., 2009). It is possible that this controversy can be settled by also studying non-facial stimuli. It therefore matters to establish first how the two kinds of information are processed when no face is present, and then to ask how the face network draws on them.

In this study, we trained participants to categorize abstract stimuli based on their configural or featural properties, and scanned them with fMRI to identify brain regions that support each type of processing. We also identified the OFA, FFA, and pSTS in individual-subjects and asked whether these regions differ in their preferences to configural or featural properties of these abstract stimuli, and whether they connect preferentially with domain-general configural and featural regions.

## Methods

### Participants

Eighteen adults (11 female; mean age = 23.9 years, SD = 5.9) completed the full experiment, a sample size within the range shown to reliably detect effects in fMRI studies of object and face processing across the ventral and dorsal visual streams (Ayzenberg & Behrmann, 2022; Bracci & Beeck, 2016; Freud et al., 2017; Jeong & Xu, 2016). Sixteen were right-handed and two were left-handed by self-report, and all had normal or corrected-to-normal vision, with those who required correction wearing MRI-safe lenses matched to their prescription throughout scanning. One additional participant was recruited but did not reach the training criterion described below and was excluded before scanning. Two participants did not complete the resting-state scan, and the connectivity analyses are based on the remaining 16. Participants were recruited from the Ohio State University community, gave written informed consent under a protocol approved by the university’s Institutional Review Board, and received monetary compensation.

### Experimental design

#### Stimuli and pre-scan training

The stimuli used to elicit configural and featural processing were taken from Zeng et al. (2026). They are simple abstract shapes constructed in the laboratory and were deliberately designed not to resemble faces or any familiar object category, so that they would not evoke domain-specific responses. The two sets are closely matched in overall appearance and differ only in the information that supports categorization (Figure 1A). Configural stimuli can be sorted correctly only by using the spatial relations among their parts, whereas featural stimuli can be sorted only by using the identity of the parts themselves, regardless of where those parts appear.

**Figure 1.**
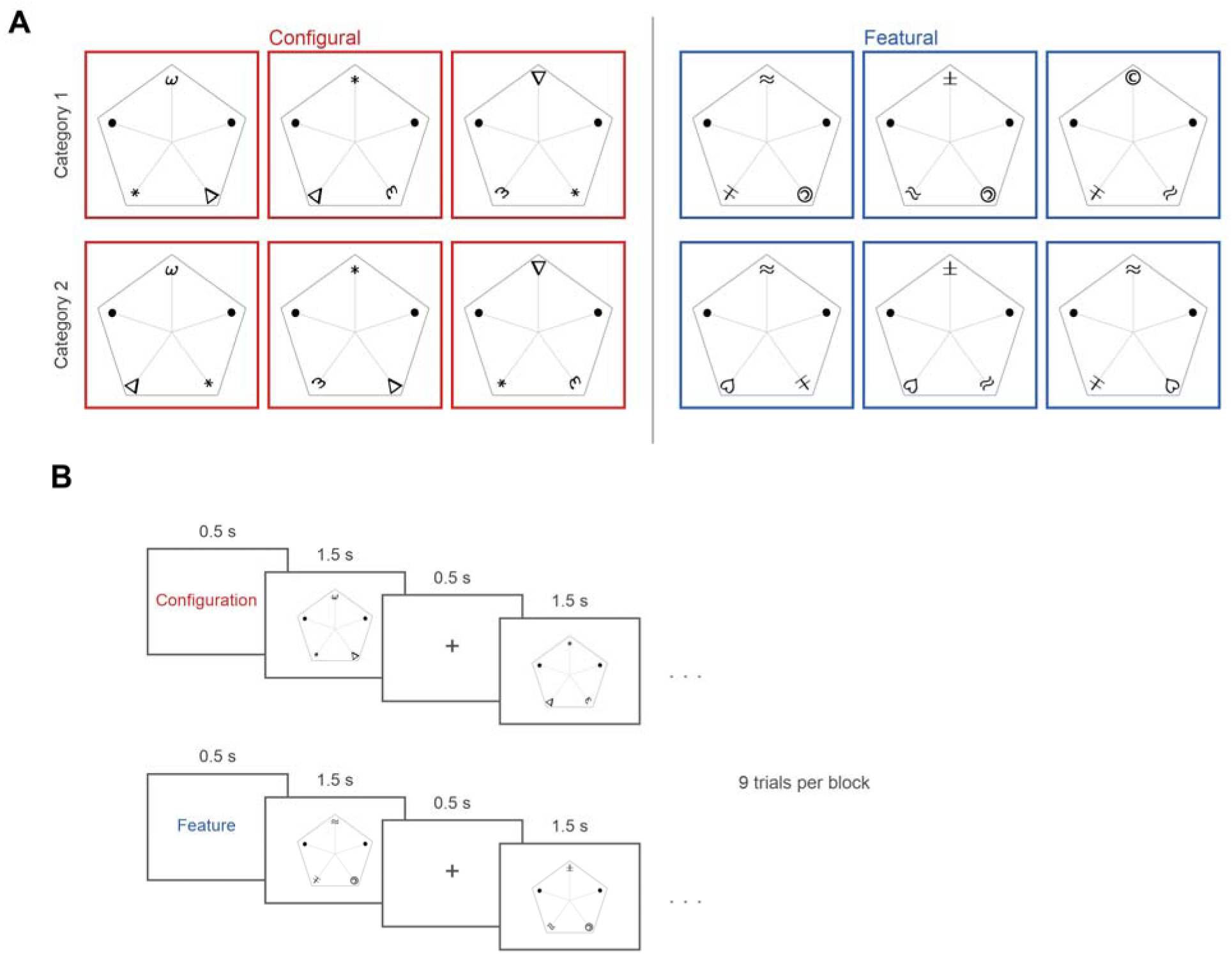
Stimuli and task. (A) Exemplars from the two stimulus sets. Within the configural set the two categories are built from the same local features and differ only in where those features sit. Within the featural set the two categories share a single layout and differ only in the features themselves. Neither set can be categorized by the property that the other set holds constant. (B) The opening trials of a configural and a featural block. Each trial ran 0.5 s of fixation followed by a 1.5 s stimulus. On the first trial of a block the fixation cross was replaced by the word Configuration or Feature, which told participants which kind of information was relevant for the nine trials of that block.

Because the two sets look so similar, participants must first learn which kind of information is relevant to each set. Each participant therefore completed a training session in the laboratory, lasting approximately 1.5 to 2 h, immediately before the scanning session on the same day. The training task was identical to that of Zeng et al. (2026) except for the number of trials and blocks. On each trial, a fixation cross appeared for approximately 0.5 s (jittered between 0.45 and 0.55 s), followed by a single stimulus that remained on the screen until the participant responded or for a maximum of 5 s. Participants sorted each stimulus into one of two groups by pressing 1 or 2, after which feedback (“Correct”, “Wrong”, or “No Response” when no key was pressed) was displayed for approximately 0.5 s (jittered over the same range). At the start of each block, a 5 s screen informed participants which kind of information was relevant in the upcoming block, that is, whether to categorize on the basis of the parts alone or on the basis of both the parts and their locations. The specific mapping between stimuli and response groups was never disclosed and had to be acquired through trial and error.

Participants completed six blocks of 150 trials in each condition, giving twelve blocks in total, presented in a fully randomized order. To verify that participants had learned to extract the relevant information, we required accuracy above 90% in both the final configural block and the final featural block, with trials on which no response was made scored as errors. This criterion was evaluated at the end of the training session, and participants who did not meet it did not proceed to scanning. Our previous work indicates that participants typically reach this level of performance within a single block, and we administered six blocks per condition so that the extraction of configural and featural information would be well practiced by the time of scanning. All 18 participants included in the analyses met this criterion.

### Main experiment

The main experiment used a block design, and the task was closely modeled on the training task. Participants categorized each stimulus into one of two groups by pressing one of two buttons on a response box held in their dominant hand. The task differed from training in three respects. First, trial timing was fixed rather than jittered. Second, each stimulus was presented for a fixed 1.5 s (by the end of training, response times were typically below 1 s). Third, no feedback was given, so that feedback-related activity would not contaminate the responses.

Each trial began with a 0.5 s fixation cross, followed by the 1.5 s stimulus, so that every trial lasted 2 s. The stimulus remained on the screen for its full duration regardless of when the response was made. Each block consisted of nine trials and therefore lasted 18 s. On the first trial of a block, the fixation cross was replaced by the word “Configuration” or “Feature”, which indicated the kind of information that was relevant for the block that followed. Within a block, stimuli were drawn at random, with replacement, from the exemplar set of the relevant condition (Figure 1B).

Each run contained 16 task blocks, eight per condition, arranged as four repetitions of an ABBA sequence, with 14 s fixation periods at the start of the run and after every fourth block. The two run orders, one beginning with a configural block and the other with a featural block, alternated across successive runs within each participant and were counterbalanced across participants. Eleven participants completed six runs and seven completed eight runs, depending on the time available in the scanning session, and all completed runs were included in the analyses.

### Face localizer

Face-selective regions were localized using the dynamic localizer from Pitcher et al. (2011a), which yields more robust and more reliably identifiable face-selective regions in individual participants than static localizers (Fox et al., 2009). Short movie clips were drawn from five categories, namely faces, bodies, scenes, objects, and scrambled objects. Each run contained two blocks per category, with six clips per block and a block duration of 18 s. Three 18 s rest blocks, during which the screen cycled through full-screen colors, occurred at the beginning, the middle, and the end of each run. The category blocks followed a palindromic order that was randomized across runs. Every participant completed two runs. To keep participants attentive, they performed a one-back task throughout, pressing a button on a response box held in their dominant hand whenever a clip repeated on consecutive presentations.

### Resting-state scan

Participants also completed a 10 min resting-state scan. A white fixation cross was presented at the center of a black screen for the duration of the scan, and participants were asked to stay awake, to keep their eyes on the cross, and not to think about anything in particular. Wakefulness was monitored throughout with an eye tracker. The resting-state scan was acquired either before or after the main experiment, depending on the scanning session.

### MRI data acquisition

All images were acquired on a Siemens MAGNETOM Prisma 3 T scanner equipped with a 32-channel phased-array head coil.

A T1-weighted anatomical volume was acquired with an MPRAGE sequence (TR = 2300 ms, TE = 2.9 ms, TI = 900 ms, flip angle = 9 degrees, 176 sagittal slices, 1.0 mm isotropic voxels, FoV = 256 × 224 mm, GRAPPA acceleration factor 2).

Functional images for the main experiment, the face localizer, and the resting-state scan were acquired with the same T2*-weighted gradient-echo echo-planar sequence (TR = 2000 ms, TE = 29 ms, flip angle = 76 degrees, 72 contiguous slices providing whole-brain coverage, 2.0 mm isotropic voxels, FoV = 216 × 216 mm, multiband acceleration factor 3, phase partial Fourier 7/8, anterior to posterior phase encoding). Each run of the main experiment comprised 179 volumes, each localizer run comprised 117 volumes, and the resting-state scan comprised 300 volumes.

### Data analysis

All analyses were carried out with FreeSurfer version 7.4.1 and its FsFast pipeline, together with custom MATLAB code (version R2024a).

### Preprocessing

Cortical surfaces were reconstructed from each participant’s T1-weighted volume using the FreeSurfer recon-all pipeline. Functional data were then processed in FsFast, with each run handled separately. Runs were motion corrected to their middle frame and registered to the participant’s own anatomical volume using boundary-based registration (Greve & Fischl, 2009). Slice timing was not corrected. The data were then resampled onto the fsaverage cortical surface and smoothed along the surface at two levels, 3 mm FWHM for the region of interest analyses and 5 mm FWHM for the whole-brain analysis. All runs were retained, and head motion was accounted for by entering the six realignment parameters as nuisance regressors in the model described below.

### First-level analysis

Responses were estimated with a general linear model implemented in FsFast, computed on the fsaverage surface and separately for each hemisphere. Each 18 s block was modeled as a single event of that duration and convolved with a gamma function (delta = 2.25 s, tau = 1.25 s). Along with one regressor per condition, the model included second-order polynomial terms to remove low-frequency drift and the six motion parameters. Runs were modeled separately and then combined within each participant. The contrast of interest was configural minus featural (C-F). Localizer data were modeled in the same way with five condition regressors, and face selectivity was defined by the contrast of Faces > Objects.

### Whole-brain group analysis

Whole-brain analyses used the data smoothed at 5 mm. For each hemisphere, the individual C-F effect sizes were concatenated across participants and submitted to a one-sample test of the group mean on the fsaverage surface. Clusters were formed at a vertex-wise threshold of p < 0.005, two-tailed, and were retained if they reached a cluster-wise threshold of p < 0.05. Cluster-wise significance was determined by precomputed Monte Carlo simulation on the fsaverage surface (Hagler et al., 2006) and Bonferroni corrected across the two hemispheres.

### Region time courses

Each participant completed the runs under two paradigm variants (ABBA and BAAB), which alternated run by run. Regions were defined from the runs of one variant and the time courses were extracted from the runs of the other. The regions were defined within search spaces built from the group-level clusters. Each cluster was dilated across the surface mesh to accommodate individual variability in the location of these regions, using 20 iterations for the configural clusters and 10 for the featural cluster. On the fsaverage surface, the parietal search space comprised 7,535 vertices in the left hemisphere and 10,265 in the right, the temporal search space 4,516 and 5,500, the frontal search space 4,320 in the left hemisphere (no frontal cluster survived correction in the right), and the featural search space 2,376 and 3,600. Within each configural search space we took the 10% of vertices with the highest C-F t values, and within the featural search space the 10% with the lowest. Percent signal change was computed relative to the mean of the fixation frames of each run and averaged across runs and then participants.

### Face-selective regions of interest

Face-selective regions were defined separately in each participant, following the group-constrained approach of Julian et al. (2012). Search spaces covered the FFA, the OFA, and the pSTS in each hemisphere. Within each search space, we took the top 10% as that participant’s region of interest. These regions were then applied to the main experiment data, which were independent of the localizer.

Responses were taken from the data smoothed at 3 mm. For each region we averaged the z statistic of the C-F contrast across its vertices. The regions were defined separately in each hemisphere, and the two hemispheres’ values were averaged in the analysis.

### Resting-state functional connectivity

Most participants were scanned with the resting-state parameters given above (TR = 2000 ms, 300 volumes). For three participants the resting-state data were taken from a separate session and were acquired with different parameters (TR = 1000 ms, 580 volumes, for two participants; TR = 5000 ms, 120 volumes, for one). Restricting the analysis to the 13 participants with the standard parameters did not change any of the results. The analyses reported below therefore include all 16 participants who completed a resting-state scan.

Resting data were motion corrected, registered to the anatomical volume, resampled onto the surface, and smoothed at 3 mm FWHM as described above. Framewise displacement was computed from the realignment parameters, and frames exceeding 0.5 mm were flagged (Power et al., 2012). These frames were first replaced by linear interpolation so that they would not distort the regression that followed (Power et al., 2014). Nuisance signals were then removed with CompCor (Behzadi et al., 2007), using the first five principal components of 2mm eroded white matter and cerebrospinal fluid masks derived from the FreeSurfer segmentation, together with 6 motion regressors and their first derivatives. The global signal was not removed, because global signal regression introduces negative correlations of ambiguous interpretation and can distort between-region comparisons of the kind reported by Murphy and Fox (2017). The flagged frames were then discarded from the residual time series, which were resampled onto the fsaverage surface. Across participants, the proportion of discarded frames ranged from 0 to 30%, and excluding the two participants who lost more than 10% of frames did not change the results.

Seed regions were derived from the clusters that survived correction in the group analysis of the C-F contrast. The configural preference appeared in superior parietal, lateral occipitotemporal, and frontal cortex. The frontal and superior parietal clusters overlap the dorsal attention network (Corbetta & Shulman, 2002; Silver & Kastner, 2009), so their coupling with the face regions could reflect attentional co-recruitment rather than the exchange of configural information. They were combined into a dorsal configural seed, reported in the supplementary material. The lateral occipitotemporal cluster lies outside this network and served as the ventral configural seed, reported in the main text. The featural seed was based on the featural preference in the fusiform gyrus. The search spaces were those described under Region time courses.

Within each search space, a participant-specific seed was defined from that participant’s C-F t-map from the main experiment, and we used the top and bottom 10^th^ percentile to define each region of interest. Any vertex a seed shared with the FFA, the OFA, or the pSTS was then removed from that seed, separately in each participant and hemisphere. There was very low overlap, and this only affected the ventral configural seed and the featural seed, removing on average 4.4 and 3.2 vertices, respectively (0.8% and 1.0% of the seed), and left the dorsal configural seed unchanged, which shared no vertex with any face region.

For each participant we extracted the mean time series of each seed and of each face-selective region, and computed connectivity as the Pearson correlation between a seed and a face region within the same hemisphere. Correlations were converted to Fisher z-values and averaged across hemispheres, giving one value per participant for each of the three seeds (dorsal configural, ventral configural, featural) and each of the three face regions.

### Statistical analysis

For the task response, the mean C-F z statistic of each region was tested against zero with a one-sample t test, separately for the FFA, the OFA, and the pSTS. A 2 (hemisphere) by 3 (region) repeated measures analysis of variance showed no interaction between hemisphere and region, so values were averaged across hemispheres.

Connectivity carried two factors and was analyzed with a 2 (seed) by 3 (region) repeated measures analysis of variance. A 2 (seed) by 3 (region) by 2 (hemisphere) analysis of variance showed no interaction involving hemisphere, so values were averaged across hemispheres. The interaction between seed and region was the term of interest, and it was followed up with paired t-tests comparing the two couplings within each region and comparing the size of that difference between each pair of regions. Because the three regions were selected a priori, these comparisons were not corrected for the number of regions.

## Results

### Behavioral performance during scanning

To ensure that participants actively processed configural and featural information while in the scanner, they continued to perform the categorization task throughout the main experiment. Accuracy was high in both conditions (configural, *M* = 92.9%, *SD* = 5.4%; featural, *M* = 93.4%, *SD* = 5.4%) and did not differ between them (*t*(17) = 0.60, *p* = 0.56). Response times showed the same pattern (configural, *M* = 0.82 s, *SD* = 0.10 s; featural, *M* = 0.83 s, *SD* = 0.10 s; *t*(17) = 0.51, *p* = 0.62). Any difference in brain activity between the conditions therefore cannot be attributed to a difference in task difficulty.

### Configural and featural information engage separate sets of cortical regions

We first asked, on a vertex wise scale across the whole cortical surface, where responses to configural stimuli differed from responses to featural stimuli. The two kinds of information were clearly dissociable across the cortex. Configural stimuli drove stronger responses than featural stimuli in bilateral superior parietal cortex and lateral occipitotemporal cortex, and at the junction of the superior frontal and precentral sulci in the left hemisphere. Featural stimuli drove stronger responses in the fusiform gyrus of each hemisphere. All clusters survived cluster-wise correction across the two hemispheres (Figure 2A, Table 1). The average time course of each set of regions further illustrates their dissociable activation patterns (Figure 2B). Using regions defined from one run of the paradigm and the time courses taken from the other runs to ensure independence, we find that the configural regions responded most during configural blocks and the featural regions most during featural blocks (Figure 2B).

**Figure 2.**
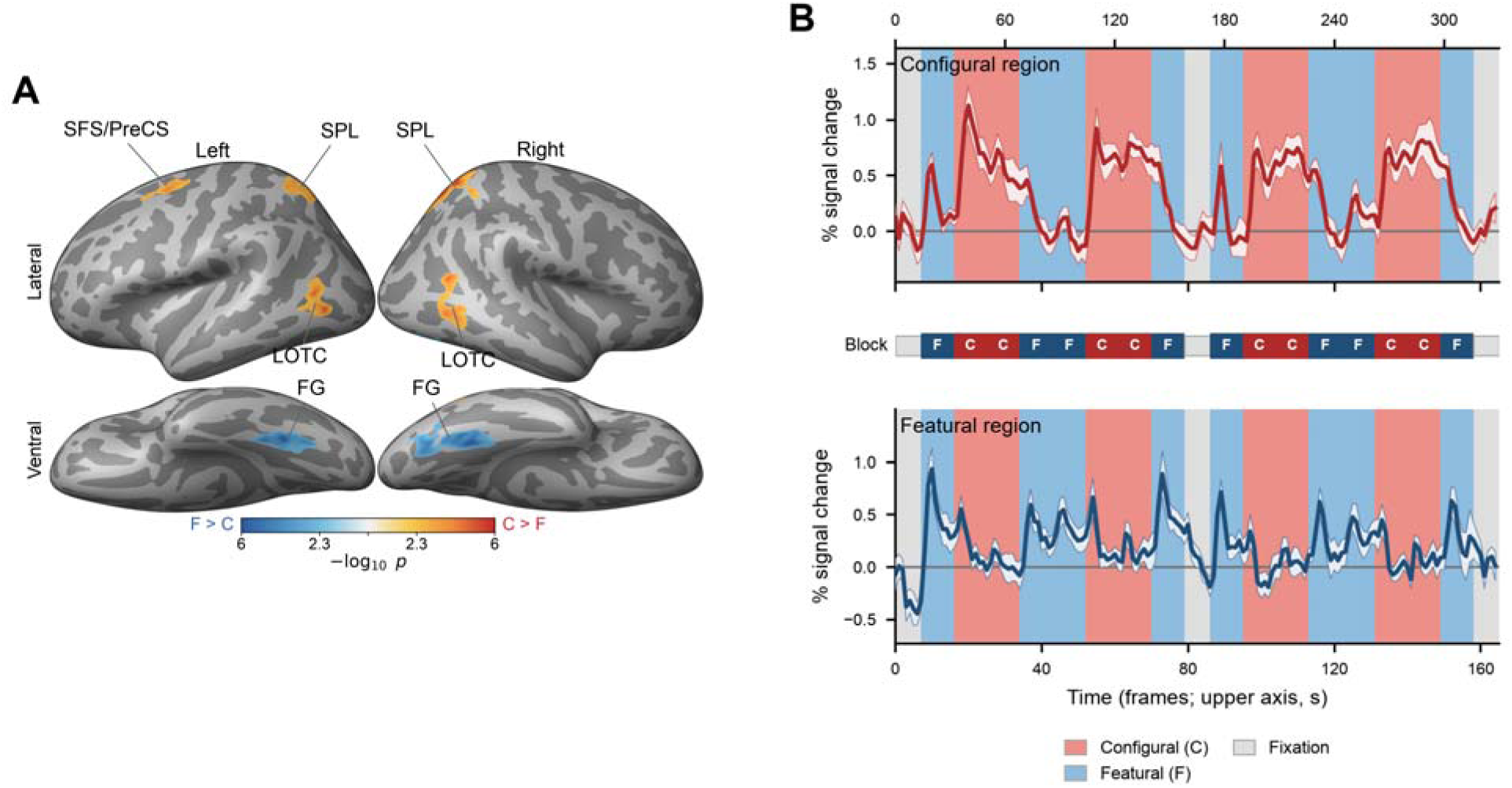
Configural and featural information engage separate sets of cortical regions. (A) Vertices where the two conditions differed, thresholded at a vertex-wise p < 0.005 and retained at a cluster-wise p < 0.05, on the inflated fsaverage surface. Warm colors mark a stronger response to the configural condition and cool colors a stronger response to the featural condition. Labeled clusters are those listed in Table 1. (B) Group mean time courses in the configural and featural regions. The regions were identified from one half of the runs and the time courses taken from the other half. Shading around the curve gives the standard error across participants, and the background marks configural (red) and featural (blue) blocks with fixation in gray.

**Table 1.** Cortical clusters showing a significant difference between configural and featural conditions (C - F) in the whole-brain surface analysis (*N* = 18).

| Hemisphere | Region | Direction | Size (mm<sup>2</sup>) | Vertices | MNI (x, y, z) | Peak $t$ | Peak $p$ | Cluster-wise $p$ |
| --- | --- | --- | --- | --- | --- | --- | --- | --- |
| Left | Superior parietal | C > F | 772.5 | 1767 | -17.6, -65.3, 51.6 | 4.80 | < 0.001 | < 0.001 |
| Left | Fusiform | F > C | 507.4 | 826 | -29.8, -54.5, -16.7 | -7.16 | < 0.001 | < 0.001 |
| Left | Lateral occipitotemporal | C > F | 377.0 | 746 | -39.9, -67.0, 0.3 | 6.26 | < 0.001 | 0.001 |
| Left | Superior frontal sulcus / precentral junction | C > F | 286.5 | 684 | -23.5, -1.3, 45.9 | 5.14 | < 0.001 | 0.008 |
| Right | Superior parietal | C > F | 1279.5 | 2776 | 22.0, -59.4, 62.6 | 6.82 | < 0.001 | < 0.001 |
| Right | Fusiform | F > C | 746.2 | 1245 | 28.1, -52.0, -15.7 | -7.25 | < 0.001 | < 0.001 |
| Right | Lateral occipitotemporal | C > F | 490.4 | 919 | 42.4, -56.9, 12.8 | 5.77 | < 0.001 | < 0.001 |

Clusters were formed at a vertex-wise threshold of $p < 0.005$ (two-tailed) and retained at a cluster-wise threshold of $p < 0.05$ , determined by Monte Carlo simulation on the fsaverage surface and Bonferroni corrected across the two hemispheres. Regions are labeled by the *aparc* parcel containing the cluster peak, except where no parcel contained a majority of the cluster's vertices. The bilateral lateral occipitotemporal clusters spanned the inferior parietal, lateral occipital, middle temporal, and inferior temporal parcels, and fell within the boundaries of lateral occipitotemporal cortex as defined by Lingnau and Downing (2015). The left frontal cluster spanned the caudal middle frontal, superior frontal, and precentral parcels in near-equal proportions and was centered on the junction of the superior frontal and precentral sulci. Coordinates give the peak vertex in MNI305 space. Peak $t$ was derived from the peak vertex-wise $p$ ( $df = 17$ ). Cluster-wise $p$ values of "< 0.001" are at the floor of the simulation distribution.

### Face-selective regions differ in their preference for configural vs. featural information

Different regions of the face network are known to serve different functions in face perception (Haxby et al., 2000; Pitcher & Ungerleider, 2021). We asked whether they also differ in their preference for configural and featural information. Because a hemisphere by region analysis of variance showed no interaction between the two hemispheres (*F*(2, 34) = 1.50, *p* = 0.24), we averaged across the two hemispheres in the analyses that follow.

We found that each face area had different preferences for featural and configural perception (Figure 3A). The pSTS preferred configural information, responding more strongly to configural than to featural stimuli (mean *z* = +0.98, *t*(17) = 3.84, *p* = 0.001, Cohen’s *d*z = 0.91). The OFA preferred featural information, responding more strongly to featural stimuli (mean *z* = -1.20, *t*(17) = -2.75, *p* = 0.014, *d*z = -0.65). The FFA sat between them and favored neither kind (mean *z* = -0.55, *t*(17) = -1.73, *p* = 0.10, *d*z = -0.41). The three regions were also distinguishable from one another. The pSTS was more configural than either the FFA (*t*(17) = 4.16, *p* < 0.001, *d*z = 0.98) or the OFA (*t*(17) = 4.36, *p* < 0.001, *d*z = 1.03), and the FFA was less featural than the OFA (*t*(17) = 2.35, *p* = 0.031, *d*z = 0.55). The two ends of this arrangement were held by regions with a preference, and its middle by a region favoring neither.

**Figure 3.**
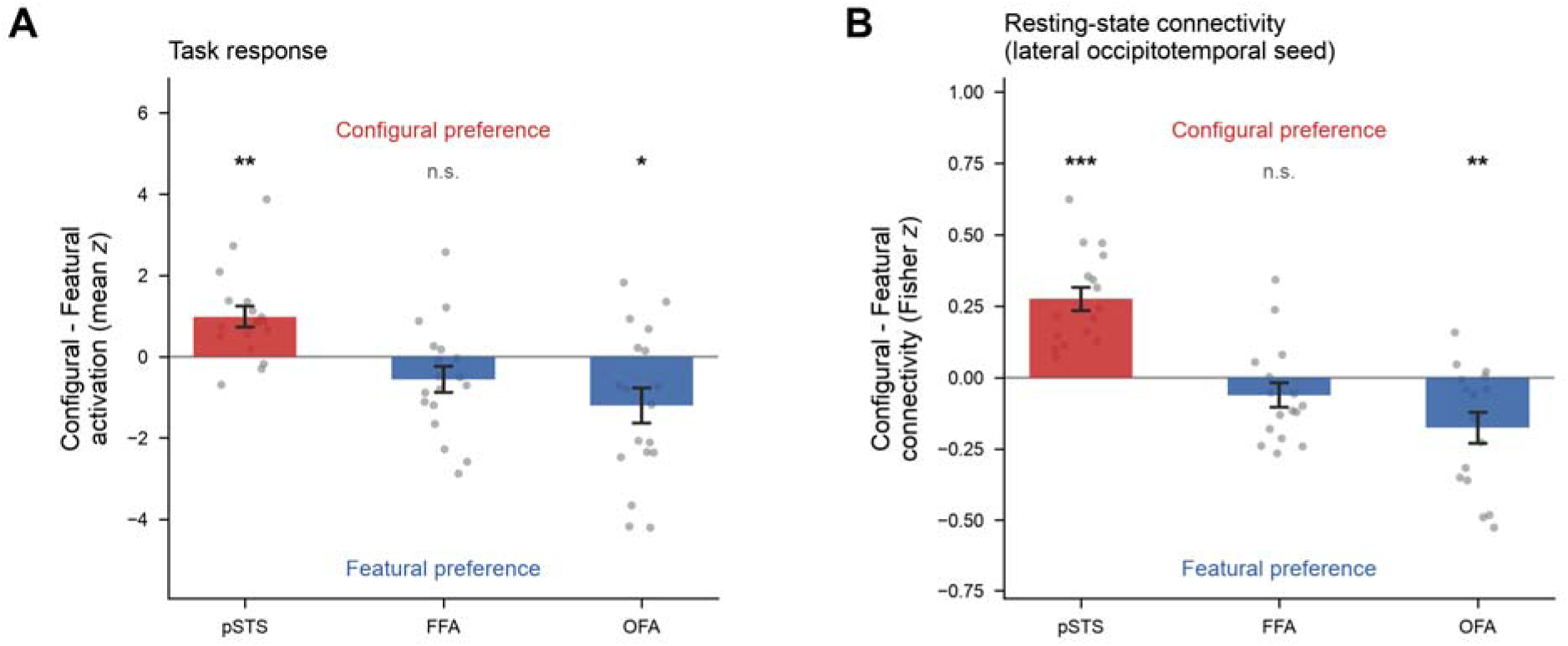
Preference for configural over featural information in the three face-selective regions. (A) Contrast between the two conditions, as the mean z statistic in each region, averaged across hemispheres. (B) Difference in resting-state coupling to the configural and the featural seed. The configural seed here is the lateral occipitotemporal cluster; the frontal and parietal clusters are shown in Supplementary Figure 1. Points are individual participants and error bars give the standard error across participants. Asterisks mark the test against zero, with *** p < 0.001, ** p < 0.01, * p < 0.05 and n.s. not significant.

### Resting-state connectivity follows the same pattern

We next asked whether the same organization is present in the brain’s intrinsic functional architecture, by measuring how strongly each face region was coupled at rest to the newly identified configural and featural regions (seed regions). Of note, these domain-general seed regions barely overlapped with the face regions, with the configural region sharing no vertex with any of them (Dice = 0) and the featural region overlapping minimally (Dice ≤ 0.012); any shared vertices were removed before the correlations were computed, such that the analysis measured coupling between distinct regions that process different information.

Because a seed by region by hemisphere analysis of variance showed no interaction involving hemisphere (all *p* > 0.10), we again averaged across the two hemispheres. We observed a seed by region interaction (*p* < 0.001; Figure 3B, Table 2; see also Supplementary Figure 1A for raw connectivity values; and Supplementary Figure 1B for dorsal configural seed regions). The pSTS was more strongly connected to the configural than to the featural seed (*p* < 0.001), the OFA showed the opposite pattern (*p* = 0.006), while the FFA fell between them, with no reliable difference in connectivity to either seed (*p* = 0.17). Post-hoc comparisons showed that the pSTS was separated from both the FFA and the OFA (both *p* < 0.001); the FFA and the OFA showed marginal differences (*p* = 0.078). This pattern of connectivity bias mirrored the activation patterns, with the pSTS at the configural end, the OFA at the featural end, and the FFA between them.

**Table 2.** Resting-state connectivity of the face-selective regions with the ventral configural and the featural seed (*N* = 16).

| Effect | *F* | df | *p* | partial $\eta^2$ |
| --- | --- | --- | --- | --- |
| Seed | 0.17 | 1, 15 | 0.683 | 0.011 |
| Region | 1.35 | 2, 30 | 0.274 | 0.083 |
| Seed × Region | 29.63 | 2, 30 | < 0.001 | 0.664 |

| Contrast | *t* | df | *p* | Cohen's *dz* |
| --- | --- | --- | --- | --- |
| pSTS, configural - featural seed | +6.76 | 15 | < 0.001 | +1.69 |
| FFA, configural - featural seed | -1.43 | 15 | 0.174 | -0.36 |
| OFA, configural - featural seed | -3.22 | 15 | 0.006 | -0.81 |
| pSTS vs FFA, difference in connectivity | +7.41 | 15 | < 0.001 | +1.85 |
| pSTS vs OFA, difference in connectivity | +6.12 | 15 | < 0.001 | +1.53 |
| FFA vs OFA, difference in connectivity | +1.89 | 15 | 0.078 | +0.47 |

## Discussion

Configural processing has been studied almost entirely using face stimuli, where a large behavioral literature has treated it as a hallmark of face perception, whether special to faces or acquired through expertise (Diamond & Carey, 1986; Farah et al., 1998; Gauthier & Tarr, 1997; Maurer et al., 2002; Tanaka & Farah, 1993). By building configural stimuli that were not faces, together with featural stimuli matched to them in task and difficulty, we were able to treat configural and featural processing as domain-general modes of extracting visual information. We found here that the two types of information are dissociable in the brain. Featural information engaged fusiform regions, falling posterior to category-selective regions, perhaps reflecting involvement in analyzing local form (Goodale & Milner, 1992; Grill-Spector & Weiner, 2014). Configural information engaged three regions, including dorsal configural clusters which are in line with supporting spatial cognition, and ventral configural clusters that lies in lateral occipitotemporal cortex of both hemispheres.

Recent work suggests that this ventral configural cluster may support sensitivity to object shape (Bracci & Beeck, 2016; Kourtzi & Kanwisher, 2000; Malach et al., 1995) and a sensitivity to the spatial relations among objects (Abassi & Papeo, 2020; Baldassano et al., 2017; Quadflieg et al., 2015; Wurm & Caramazza, 2019), which support our current findings. Configural information consists of exactly these two together, the shapes of the parts and their arrangement, which is what distinguishes it from featural information. In this prior literature, however, the relations were always carried by people and the objects they act on, or by one person and another, and the accounts built on them followed from the stimuli, placing this cortex either in a pathway for action recognition or in a network for social perception (Pitcher & Ungerleider, 2021; Wurm & Caramazza, 2022). Our stimuli contained no bodies, no actions, and no category information of any kind, and the preference for configural information appeared there nonetheless, so our findings suggest that the sensitivity of this cortex to spatial relations may not depend on action or social content. It is possible that action and social perception recruit the region because they themselves depend on configural information; i.e., since body posture, the relative orientation of two people, and object-directed actions are each defined by parts and their spatial arrangement. Thus it is possible that this cortex computes domain-general configural information, and action and sociality are domains that simply draw on this processing.

Having isolated the two kinds of information in this way, we could then ask how the face network, which relies on both kinds of information, responds and interacts with these regions. We find, using both task activation and resting-state connectivity, that the pSTS lies on the configural end, the OFA at the featural end, and the FFA between the two. This result is in line with prior literature. The OFA is thought to lie at the earliest stage of the face network and is generally treated as its entry point, computing a part-based description that is passed forward to later regions (Haxby et al., 2000; Pitcher et al., 2011b). What it codes is specifically local. Disrupting the right OFA with TMS impairs the discrimination of individual face parts but leaves the spacing between them intact (Pitcher et al., 2007), and the region tracks the presence of parts rather than their configuration (Liu et al., 2010). Reading out local parts is in essence featural processing, which places the OFA at the featural end.

The pSTS is on the opposite end. It is tuned to the changeable aspects of faces, responding to moving eyes and mouths, to expression and gaze, and more broadly to the social signals a face conveys (Pitcher & Ungerleider, 2021; Puce et al., 1998). These signals are carried largely by configuration, since a change of viewpoint alters the distances among the features and a change of expression rearranges them, as when the brows draw together in anger. Perceiving them therefore depends on configural information (Calder et al., 2000), placing the pSTS at the configural end.

The FFA ran against our expectation. Because face identification is usually attributed to configural processing, and because configural tasks with non-face objects of expertise engage this region (Gauthier et al., 1999), we expected it to respond more to configural than featural processing. Instead, it did not favor either configural or featural information and additionally showed no differentiation in its connectivity with configural or featural regions. This result is in line with theories suggesting that the FFA is usually positioned downstream of the OFA, integrating part-based input into a unified representation of a face and its identity (Freiwald & Tsao, 2010; Haxby et al., 2000), a computation that needs both the parts and the configuration they form.

There exist several limitations in this study. First, we used a block design in our experiment in order to obtain a high signal-to-noise ratio (Birn et al., 2002; Friston et al., 1999). This favored sensitivity to the contrast between configural and featural processing, but it collapsed the finer, trial-level structure that our stimuli carried, including the differences among individual exemplars and among categories. The data therefore cannot support multivariate analyses such as representational similarity analysis or decoding (Haxby et al., 2001; Kriegeskorte et al., 2008; Norman et al., 2006), which would characterize how each kind of information is represented within a region. An event-related design would preserve that trial-level variation. The second limitation concerns connectivity. Being correlational, it shows that the face regions are coupled to the configural and featural systems but not the direction in which information flows (Friston, 2011). Our proposal that the two act as mid-level signals feeding forward into high-level, category-selective cortex (DiCarlo et al., 2012; Felleman & Van Essen, 1991) therefore awaits evidence sensitive to directionality, which effective-connectivity modeling or causal manipulation could provide (Friston et al., 2003; Zachariou et al., 2017).

## Supplementary material

## Supplementary material

**Supplementary Figure 1.**
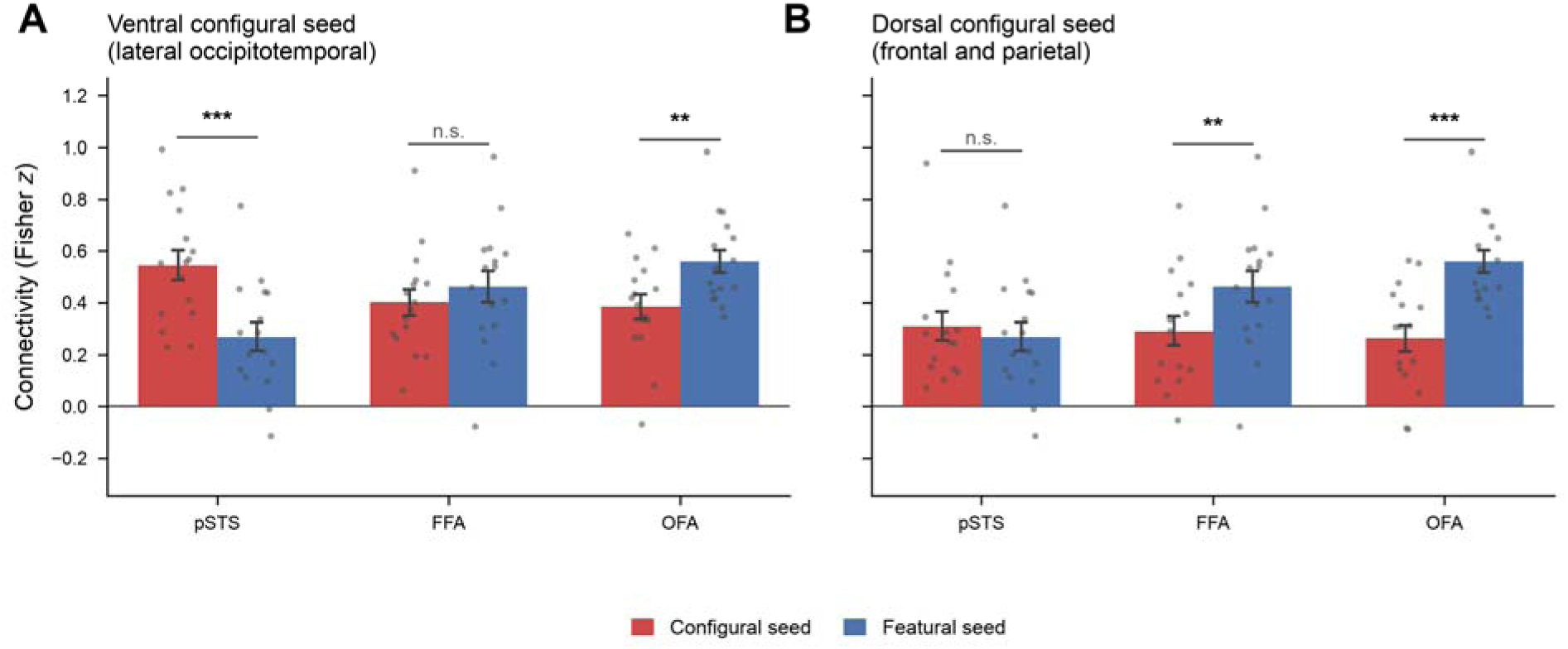
Raw functional connectivity values behind the differences in Figure 3B, for each seed and each face region, in Fisher z (N = 16). (A) The ventral configural seed and the featural seed. (B) The dorsal configural seed, combining the frontal and parietal clusters, and the featural seed. The mark above each region indicates a significant between pairs. For the dorsal seed, connectivity was lower than to the featural seed in the FFA (z = -0.171, t(15) = -3.65, p = 0.002) and the OFA (z = -0.295, t(15) = -5.27, p < 0.001), and did not differ in the pSTS (z = 0.040, t(15) = 1.12, p = 0.28). Points are individual participants and error bars give the standard error across participants, with *** p < 0.001, ** p < 0.01 and n.s. not significant.

